# The evolutionary advantage of heritable phenotypic heterogeneity

**DOI:** 10.1101/028795

**Authors:** Oana Carja, Joshua B. Plotkin

## Abstract

Phenotypic plasticity is an evolutionary driving force in diverse biological processes, including the adaptive immune system, the development of neoplasms, and the bacterial acquisition of drug resistance. It is essential, therefore, to understand the evolutionary advantage of an allele that confers cells the ability to express a range of phenotypes. Of particular importance is to understand how this advantage of phenotypic plasticity depends on the degree of heritability of non-genetically encoded phenotypes between generations, which can induce irreversible evolutionary changes in the population. Here, we study the fate of a new mutation that allows the expression of multiple phenotypic states, introduced into a finite population otherwise composed of individuals who can express only a single phenotype. We analyze the fixation probability of such an allele as a function of the strength of inter-generational phenotypic heritability, called memory, the variance of expressible phenotypes, the rate of environmental changes, and the population size. We find that the fate of a phenotypically plastic allele depends fundamentally on the environmental regime. In a constant environment, the fixation probability of a plastic allele always increases with the degree of phenotypic memory. In periodically fluctuating environments, by contrast, there is an optimum phenotypic memory that maximizes the probability of the plastic allele’s fixation. This same optimum value of phenotypic memory also maximizes geometric mean fitness, in steady state. We interpret these results in the context of previous studies in an infinite-population framework. We also discuss the implications of our results for the design of therapies that can overcome resistance, in a variety of diseases.

## Introduction

Perpetual volatility in the surrounding microenvironment, nutrient availability, temperature, immune surveillance, antibiotics or other drugs are realities of life as a microorganism [1; 2; 3]. To persist in constantly changing environments and increase resilience, microbial populations often employ mechanisms that expand the range of phenotypes that can be expressed by a given genotype [3; 4]. This form of bet-hedging may not confer an immediate fitness benefit to any one individual, but it can sometimes act to increase the long-term survival and growth of an entire lineage [5; 6].

Phenotypic heterogeneity has been well documented both in the context of cellular noise without any known environmental triggers [7] and in the context of persistent environmental challenges [8]. Classic examples include the bifurcation of a genotypically monomorphic population into two phenotypically distinct bistable subpopulations [1], or phase variation, a reversible switch between different phenotypic states driven by differences in gene expression [3; 4; 9]. In order to motivate our modeling study, we first describe and compare four clinically relevant examples of phenotypic bet-hedging and its role in pathogen survival and the evolution of resistance.

One of the most striking examples of evolutionary bet-hedging is bacterial persistence [6; 10; 11], whereby a genetically monomorphic bacterial population survives periods of large antibiotic concentrations by producing phenotypically heterogeneous sub-populations, some of which are drug-insensitive [12]. Persister phenotypes constitute dormant, transient phenotypic (epi)states, protected from the action of drugs. Phenotypic variants that can survive treatment differ from traditional resistance mutants, because the dormant phenotypes do not rely on genetic mutations [12]. The dormant phenotypic state can be partially heritable upon cellular division, so that the offspring cell can “remember” and express the phenotypic state of its parent with a certain probability. In such bacterial populations individuals typically acquire and relinquish the dormant phenotype at rates much higher than that of DNA mutation [13], providing the population with the phenotypic plasticity needed to persist through periodic environmental stresses. Even though persisters are a non-genetic form of inheritance, the capacity to generate persistent cells, and the propensity to retain the phenotype of a parent, are likely under genetic control [14].

There are surprising parallels between antibiotic persistence in bacteria and the quiescent phenotype in cancer cell populations [15]. Although genetic resistance alleles and their role in tumor dynamics have been a focus in cancer research, there has been less study of drug-tolerant epi-phenotypes and their impact on the evolution of a neoplasm [16]. Under drug concentrations that can eradicate some cancer cells, these epiphenotypes can preserve viability by producing genetically identical, but slower-dividing sub-populations. The slower-dividing cells are maintained in the population by their ability to revert to the faster-growing phenotype in the absence of drug pressure. The drug-torelant epi-state can be acquired, inherited, and relinquished by cells at rates much higher than that of genetic mutation and, similar to antibiotic persisters, constitutes a mechanism by which protected subpopulations of cells can escape periods of high drug concentrations. Just like bacterial persisters, this bet-hedging strategy confers on the population a degree of phenotypic heterogeneity that helps it withstand periods of environmental stress.

Neoplastic cell populations employ other bet-hedging strategies during growth and spread. Depending on mechanical cues in the tumor microenvironment, for example, populations of cancer cells can utilize heterogeneity in the fast- or slow-locomotion phenotype, producing radically different evolutionary dynamics of cancer cell populations [17]. Genetically identical cells can express and switch between different motility states. Changing the tumor microenvironmental cues alters the rate of switching between the slow-locomotion mode and the fast-locomotion mode, and therefore changes the number of cells expressing either motility phenotype [17].

Environmental shifts also occur when pathogens are transmitted between hosts. Bet-hedging in this context can manifest as a form of life-history strategy [18; 19]. Viral latency is a heterogeneous fate decision that can help a viral population persist and adapt as it encounters different host immune environments. Examples include the choice between HIV latency or active replication upon infecting a CD4+ T lymphocyte [18; 20], or latency in herpes viruses [21]. A probabilistic switch that produces both infected cells and latency with the possibility of later infectivity may confer a competitive advantage in a fluctuating environment, driven by differences in the numbers of susceptible hosts or variation in host immune system repertoires over time.

The tradeoffs of phenotypic variability in viral latency, or neoplastic motility are similar to those for antibiotic persisters or quiescent cancer cells: decreased environmental uncertainty versus decreased average replication rates. There are striking similarities among the four examples presented above: all involve genetically identical populations, with two or more available phenotypes, with each phenotype beneficial in a different environmental state. Phenotypic states are partly heritable by offspring cells. And yet individuals can switch between phenotypic states at rates that greatly exceed those of genetic mutations. Finally, the rate of ‘phenotypic mutation’ is itself under genetic control. These types of epigenetic bet-hedging strategies by dynamic regulation of phenotypic variability may allow the persistence of a population until more permanent genetic strategies can be found [22; 23]. Although the idea of phenotypic plasticity, and even subsequent genetic assimilation, dates back to Waddington [24], its critical role in microbial population dynamics has only recently been appreciated [25; 26; 27].

The ubiquity of heritable phenotypic heterogeneity demands rigorous study of its evolutionary dynamics in a population-genetic framework. Here we study the evolutionary fate of an allele that permits the expression of multiple phenotypic states, in a finite population of fixed size under either constant or periodically fluctuating environments. We imagine these phenotypic states as partially heritable, so that an expressed phenotype will be inherited by the offspring with some probability, *p*. We call this probability *p* the phenotypic memory. We study the probability of fixation of a new mutation that permits this form of heritable, increased variance in the expressed phenotype, in a population that is otherwise composed of individuals that are unable to express variable phenotypes. We are especially interested in the fate of such a mutation as a function of the variance in expressible phenotypes that it confers, the extent of phenotypic memory between generations, the rate of environmental change, and the population size. We will show that, in constant environments, increasing phenotypic memory always increases the fixation probability of such a phenotypically-variable mutant. In contrast, in periodically changing environments there is an intermediate rate of phenotypic memory that maximizes the fixation probability of such a new mutant. Moreover, the phenotypic memory that maximizes fixation probability is inversely proportional to the rate of environmental change, and it is largely insensitive to the population size or to the expressible phenotypic variance of the mutant type. Furthermore, we show that the phenotypic memory that maximizes invasion probability also maximizes geometric mean fitness in mutation-selection-drift stationary state, and we discuss these results in the context of prior studies in an infinite population.

## Evolutionary bet-hedging models

### Previous theory

Phenotypic variability is often caused by switches between different regulatory states that produce bi-or multi-stability, due to fluctuations in levels of methylation at CpG sites, for example in mRNA transcription, or protein translation [28]. Our focus here, however, is not on the specific mechanisms or bio-physical forces that govern these phenotypic state transitions [9; 19; 29]. Instead, we aim to elucidate the population-level consequences of increased phenotypic availability from a given genotype and how this non-genetic variability determines the long-term evolutionary outlook of a population [30; 31]. In this framework, the trade-offs between the evolutionary advantages of phenotypic variability and the costs of maladaptation is our principal object of inquiry. Mathematical models that incorporates stochastic fluctuations in selection pressure are already essential components of population-genetic theory [32; 33; 34; 35]. Early works by Gillespie [36; 37] established the geometric mean fitness principle, and showed that the evolution of a population under fluctuating selection is controlled not only by the mean fitness in a generation but also by the variance in fitness between generations, so that an allele with higher mean fitness can be outcompeted by one with lower mean fitness if the variance of the latter is sufficiently lower than of the former. In subsequent work, Gillespie also showed, for variation in fitness within a generation, that the strength of selection for reduced variance is inversely proportional to the population size [38; 39; 40; 41].

In all the models of variable fitness summarized above, the phenotype of an offspring is independent of the phenotype of its parent. The absence of phenotypic inheritance in these previous models make them a poor choice for studying the types of phenotypic heterogeneity described in our Introduction. Heritability of phenotype is an essential requirement for evolutionary change to occur, and so (partial) heritability of an individual’s phenotype is an essential component of the model we develop here. This form of heritable phenotypic variation we study, which produces phenotypic variability with familial correlations, is an intermediate between genetically determined phenotypic variation, where offspring are phenotypically similar to parents, and phenotypic plasticity, which causes large phenotypic variation within a genetically clonal population but is not inherited by offspring.

### A population-genetic model

We use a Wright-Fisher-type model to describe changes in allele frequencies in a finite population of fixed size *N*. Each individual is defined by one biallelic locus *A/a,* which controls its phenotype. The *A* allele encodes a fixed phenotypic value, whereas individuals with the *a* allele may express a wider range of phenotypes: such an individual’s phenotype is drawn from a random variable with a fixed mean and variance. The phenotypic random variable associated with allele *a* can either be discrete (e.g. discrete epi-phenotypes around a fixed value), or continuous (e.g. a continuous phenotypic interval available to the persister genotype *a*). We consider a population initially fixed for the wild-type allele *A*, which expresses only a single phenotype. We introduce one copy of the *a* allele in the population and determine the fixation probability of this new mutation, which confers its holders with access to a larger, and partially heritable, phenotypic range.

We study two versions of the model, one for a constant environment and one for a periodic environment (**Figure 1, A** and **B**). The mapping from phenotype to fitness depends on the environmental regime, as presented below.

**Figure 1:**
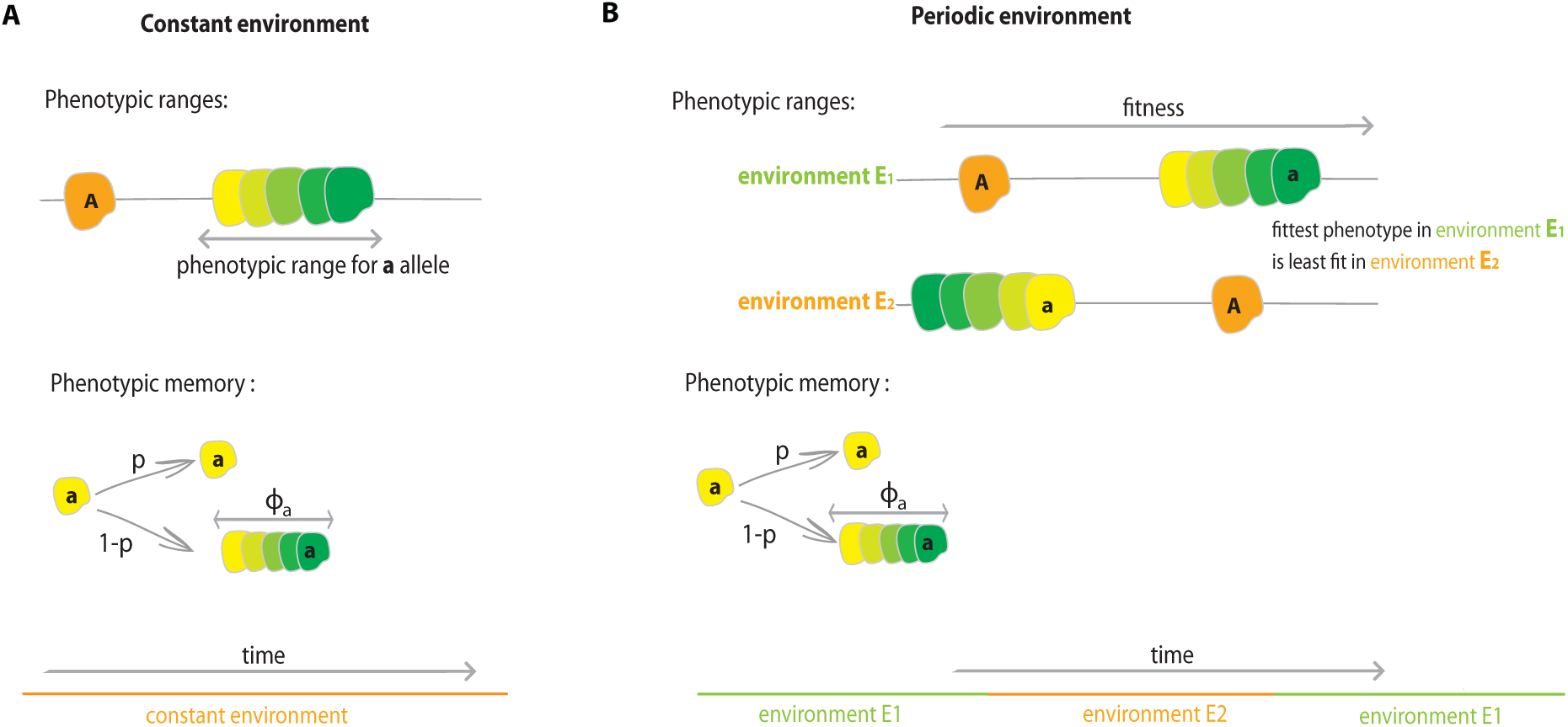
Illustration of model. **A: Constant environment.** The population is initially fixed on allele *A*, which can express only one phenotype, Φ*_A_*. We determine the fate of a new mutant, *a*, which has access to a wider phenotypic range described by the random variable Φ*_a_*. Fitness is defined to equal phenotype, and both alleles have the same mean fitness: 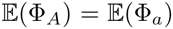. For individuals of *a* genotype there exists a probability of phenotypic memory *p*, between parent and offspring: with probability *p*, the offspring keeps the same phenotypic state as its parent, while, with probability 1 − *p*, the offspring’s phenotype is determined as a new sample from Φ*_a_*. **B: Periodic environment.** With environments changing, we assume that the phenotypes that are fitter in one environment are less fit in the other. Both alleles have the same mean fitness in their preferred environment: 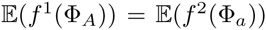 and 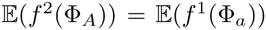 and the phenotypic variance of the *a* allele is the same in both environments: 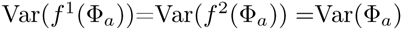.

### Constant environment

In a constant environment the fitness scheme takes the form

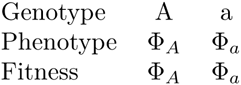

where Φ*_A_* and Φ*_a_* are random variables. The fitness of an individual is defined to be equal to its phenotype. The random variable Φ*_a_* is in fact deterministic with variance zero – that is, an individual with the *A* allele can express only one phenotype. By contrast, Φ*_a_* is not deterministic and it can be either a discrete or continuous random variable with positive variance. Φ*_A_* and Φ*_a_* are chosen so that two alleles have the same mean fitness: 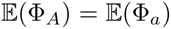. We choose fitness functions with equal means so as to focus our analysis on the effect of variance in phenotypes expressed by allele *a*, Var(Φ*_a_*), and not on any mean-fitness effect. An illustration of this model is presented in **Figure 1, Panel A**.

### Periodic environment

We study the evolution of bet-hedging in the presence of periodic fluctuations in fitness regimes. We assume that the population experiences two different environments, *E*_1_ and *E*_2_, which alternate deterministically every *n* generations, so that the both environments are experienced every *2n* generations. We assume that one environment is more favorable on average to one allele, and the other environment to the other allele.

The fitness scheme can be written as

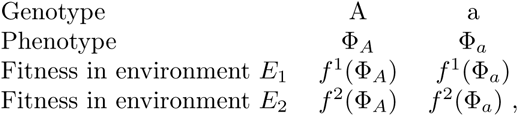

where Φ*_A_* and Φ*_a_* are random variables and the random variable Φ*_A_* is, again, deterministic with zero variance. The functions *f^i^*: ℝ→ ℝ (*i ∈* {1, 2}) map phenotype to fitness in each of the two environments, and *f*^1^ is the identity function. We assume that both alleles have the same mean fitness in their preferred environment, and the same mean fitness in their unpreferred environment: 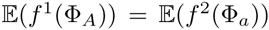 and 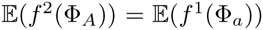. This condition also ensures that the average of two alleles’ mean fitnesses, which we denote 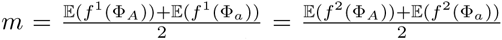, is the same in both environments. The function *f*^2^ is defined as a reflection of *f*^1^ around *m*: for any phenotype *x*, *f*^2^(*x*) = 2*m* − *f*^1^(*x*). As a result, the variance in fitness of allele *a* is the same in both environments: Var(*f*^1^(Φ*_a_*))=Var(*f*^2^(Φ*_a_*)) = Var(Φ*_a_*). These fitness functions describe a model in which each genotype has one preferred environment, but allele *a* can express a range of phenotypes whereas allele *A* expresses only a single phenotype in each environment (see illustration in **Figure 1, Panel B**). The symmetry conditions we have imposed on mean fitnesses allow us to focus our analysis on the effects of phenotypic and fitness variation.

At each generation the population experiences selection and reproduction, followed by a possible phenotypic change for offspring of allelic type *a*. In particular, to create the next generation we choose *N* individuals to reproduce from the current population, sampled with replacement with probabilities proportional to their fitness relative to the population mean fitness. We then determine the phenotypic state of every offspring in the next generation as follows. If the individual chosen to reproduce has genotype *A*, then the phenotypic state of the offspring always equals its parent’s phenotypic value. For individuals of *a* genotype, however, there exists a probability of phenotypic memory, denoted by the parameter *p*, between parent and offspring: with probability *p* the offspring retains the phenotypic state of its parent, and with probability 1 − *p* the offspring’s phenotype is drawn independently from the random variable Φ*_a_*. Thus, individuals of type *a* can express a range of phenotypic values, and their phenotype is partly heritable between generations (provided *p* > 0).

We study the possible long-term advantage of the phenotypic plasticity by analyzing the fixation probability of a phenotypically variable *a* allele introduced into a population otherwise composed of the non-variable *A* allele. How does this probability of fixation depend on environmental factors, such as the environmental period 2*n*, on demographic factors, such as the population size *N*, and on molecular factors, such as the variance in phenotypes that can be expressed by *a,* Var(Φ*_a_*), or the degree of phenotypic memory, *p*.

Although part of our analysis is analytical, we primarily determine the probability of fixation in changing environments by Monte Carlo simulation. We estimate this probability by simulating an ensemble of 10,000 replicate populations. An ensemble this large allows us to reject the null hypothesis of a neutral fixation probability with power 0.8, and it allows us to distinguish between fixation probabilities as similar as 0.001 and 0.002. We initiate all populations with one copy of the *a* allele, and we simulate the process until fixation of one of the two alleles, or discard simulations that have not absorbed after 10,000 generations (a conservative cutoff, leading to extremely few discards see **Supplementary Figure S1**).

## Results

### Constant environment results

In a constant environment the probability that an *a* allele introduced in one copy will eventually fix in the population is an increasing function of phenotypic memory, *p* (**Figure 2**). For larger values of phenotypic memory, the selective advantage of the phenotypically heterogeneous allele is increased, and this increase is more pronounced when the *a* allele can express a greater diversity of phenotypes, i.e. for Var(Φ*_a_*) large. There is a simple intuition which explains these results. The high-fitness variants of the *a* allele are preferentially transmitted to the next generation, giving the *a*-lineage an advantage; and the phenotypic memory *p* gives such individuals the opportunity to remain high-fitness instead of being resampled from the whole distribution Φ*_a_*. As a result, the mean fitness within the *a*-allele subpopulation is increased by phenotypic memory, while the variance within the *a* subpopulation is decreased. According to this simple intuition the fixation probability of the phenotypically variable *a* allele will always be greater when a alleles can express a broader range of phenotypes, that is larger Var(Φ*_a_*), as confirmed in **Figure 2**.

**Figure 2:**
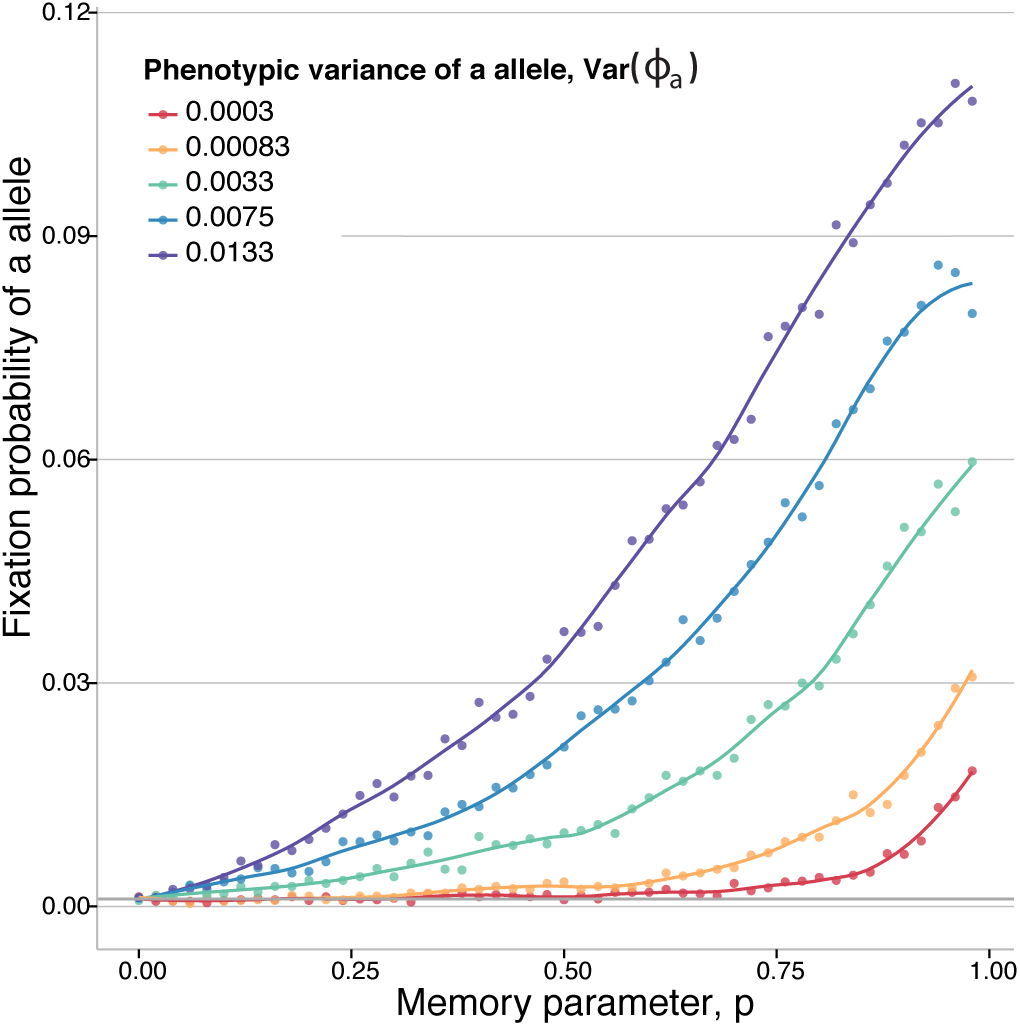
Fixation probability of a phenotypically variable allele in a constant environment. The phenotype of the *A* allele si fixed at Φ*_A_* = δ_0.8_. The distribution of phenotypes Φ*_a_* that can be expressed by the *a* allele is assumed uniform with mean also equal to 0.8. Colors represent different phenotypic variances, with Var(Φ*_a_*) presented in the legend. The dots show the fixation probabilty of a new *a* allele in a population of size *N* = 1000, as a function of the phenotypic memory parameter *p*, as determined by Monte Carlo simulation. The grey line is drawn at 1/*N*. Curves show cubic splines fitted to the simulated data.

In the absence of memory (*p* = 0), we recapture a model studied by Gillespie [38; 39] in which he found a fitness cost associated with within-generation phenotypic variance. However, the size of this fitness cost is extremely small in general, and so small in the regimes we study that the fixation probability of the *a* allele is not statistically different from 1/*N*, with power 0.8.

Our result on the advantage of phenotypic memory depends critically on individual-based memory and phenotypes. In other words, in our model, each offspring of an *a*-type parent independently chooses whether to retain its parent’s phenotype, and, if not, draws a new phenotype independently from all other offspring. Our results do not hold under an alternative model that assumes “global” choices for inheritance of phenotypes. Under that form of global sampling, all *a*-type offspring inherit their parental phenotype with probability *p*, or they simultaneously redraw the same new phenotype with probability (1 − *p*). Thus, under global phenotypic memory, all individuals of allelic type *a* express the same phenotype in each generation, similar to a model of between-generation phenotypic variance studied by Gillespie [36]. This form of global phenotypic resampling does not confer an advantage to high-fitness phenotypic realizations of the *a* allele, because low-fitness individuals are just as likely to express and retain their phenotype as high-fitness individuals (**Supplementary Figure S2**). And so, in summary, we find that phenotypic memory provides an advantage to a plastic allele *a* only when individuals independently inherit or re-sample phenotypes.

Aside from the Monte Carlo simulations shown in **Figure 2**, we can also derive an analytic approximation for the probability of fixation of the *a* allele, in the more tractable case of a discrete random variable Φ*_a_* with two available phenotypes, Φ*_a_,_min_* and Φ*_a_,_max_* (as illustrated in **Supplementary Figure S3**). The probability of fixation of the *a* allele is invariant to the choice of distribution (continuous or discrete), provided the distributions have equal variance (**Supplementary Figure S4, Panels A** and **B**). To derive an analytical approximation, we assume that the *a* mutation is first introduced with phenotype Φ*_a_,_max_* and we compute an effective selection coefficient for the entire, phenotypically variable *a* lineage. To do so we assume that the two phenotypes within the *a* lineage are immediately in mutation-selection balance. That is, given “mutation” rate 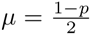 between the two phenotypes, at equilibrium, the frequency of phenotype Φ*_a,max_* within the *a*-lineage is given by *f_a,max_*:

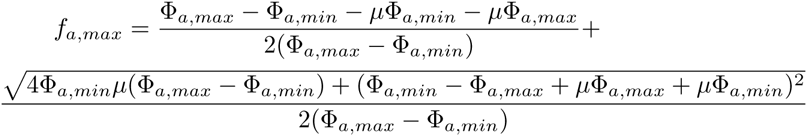

We then compute the effective selective coefficient of the *a* lineage compared to the *A* lineage as

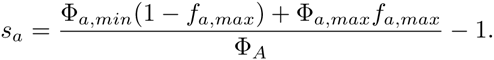

Finally, using the classic result of Kimura [42; 43], the fixation probability of the *a* allele can be approximated by 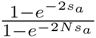. This approximation compares favorably to Monte Carlo simulations (**Supplementary Figure S4, Panel C**).

### Periodic environment results

The role of phenotypic memory is fundamentally different in a periodic environment as opposed to a constant environment. Whereas in a constant environment greater phenotypic memory always increases the advantage of the phenotypically variable *a* allele, we find qualitatively different behavior in a periodically changing environment: the fixation probability of the *a* allele is no longer monotonically increasing with phenotypic memory, but rather there exists an intermediate memory *p\** that maximizes the fixation probability.

The fixation probability of the plastic *a* allele is shown in **Figure 3, Panel A** as function of the strength of phenotypic memory *p*. The non-monotonicity we observe makes intuitive sense. One the one hand, it is beneficial to the *a* allele to have at least some phenotypic memory within each environment (*E*_1_ or *E*_2_), for the reasons described above in the case of a constant environment. But on the other hand, too much phenotypic memory is deleterious in a periodically changing environmental regime, because once the environment shifts, the *a* lineage would benefit from changing its phenotype to the new optimum. According to this simple intuition, the optimal phenotypic memory *p\** should be smaller when the environment fluctuates more rapidly, as can be seen in **Figure 3, Panel A** (comparing, say, environmental duration *n* = 5 generations to *n* = 30 generations). Therefore, the maximum phenotypic memory *p\** beyond which further memory is deleterious, should scale inversely with the environmental duration, as confirmed in **Figure 3, Panel B**. The same figure shows that the value of *p^*^* that maximizes the fixation probably of the *a* allele is insensitive to variation in the population size *N*, at least over the order of magnitude variation that we investigated.

**Figure 3:**
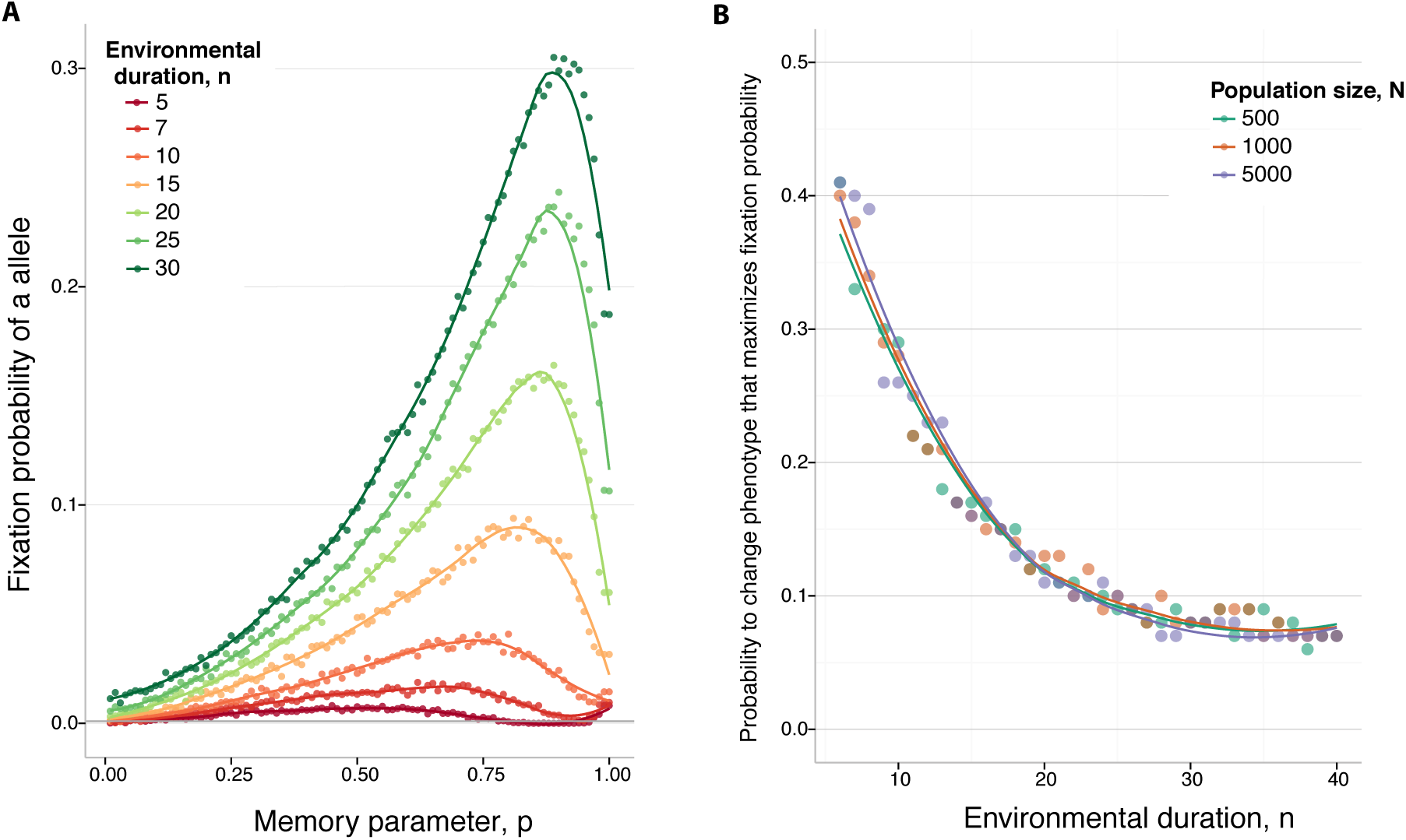
Fixation of a phenotypically variable allele in a periodic environment. **Panel A:** The dots show the fixation probability of a novel *a* allele in a population of size *N* = 5000 as a function of the phenotypic memory parameter *p*, determined by Monte Carlo simulation. In all cases we set *f*^1^(Φ*_A_*)) = 0.6 and *f*^2^(Φ*_A_*)) = 0.8. The fitness of an *a* allele in the first environment, *f*^1^(Φ*_a_*) is sampled uniformly in the interval [0.7,0.9], wheres *f*^2^(Φ*_a_*) is uniform on the interval [0.5, 0.7]; and so that each allele as the same expected fitness in its preferred environment. The initial environment is *E*_1_. The colors represent different rates of environmental change, as presented in the legend. **Panel B**: The environmental duration is plotted on the *x*-axis against the value of phenotypic memory, *p*,* that maximizes the probability of fixation of a novel *a* allele. Different colors represent different population sizes, *N*, as indicated. The phenotypic variance of the *a* allele, Var(Φ*_a_*), is set to 0.0133, and *N* = 5000. The curves show cubic spline fits to the simulated data.

Regardless of the strength of phenotypic memory, the fixation probability of a new *a* allele always increases with the variance in phenotypes it can express, Var(Φ*_a_*) (**Figure 4, Panel A**). Nonetheless, the optimum phenotypic memory for invasion, *p*,* is insensitive to the phenotypic variance of the *a* allele (**Figure 4, Panel B**).

**Figure 4:**
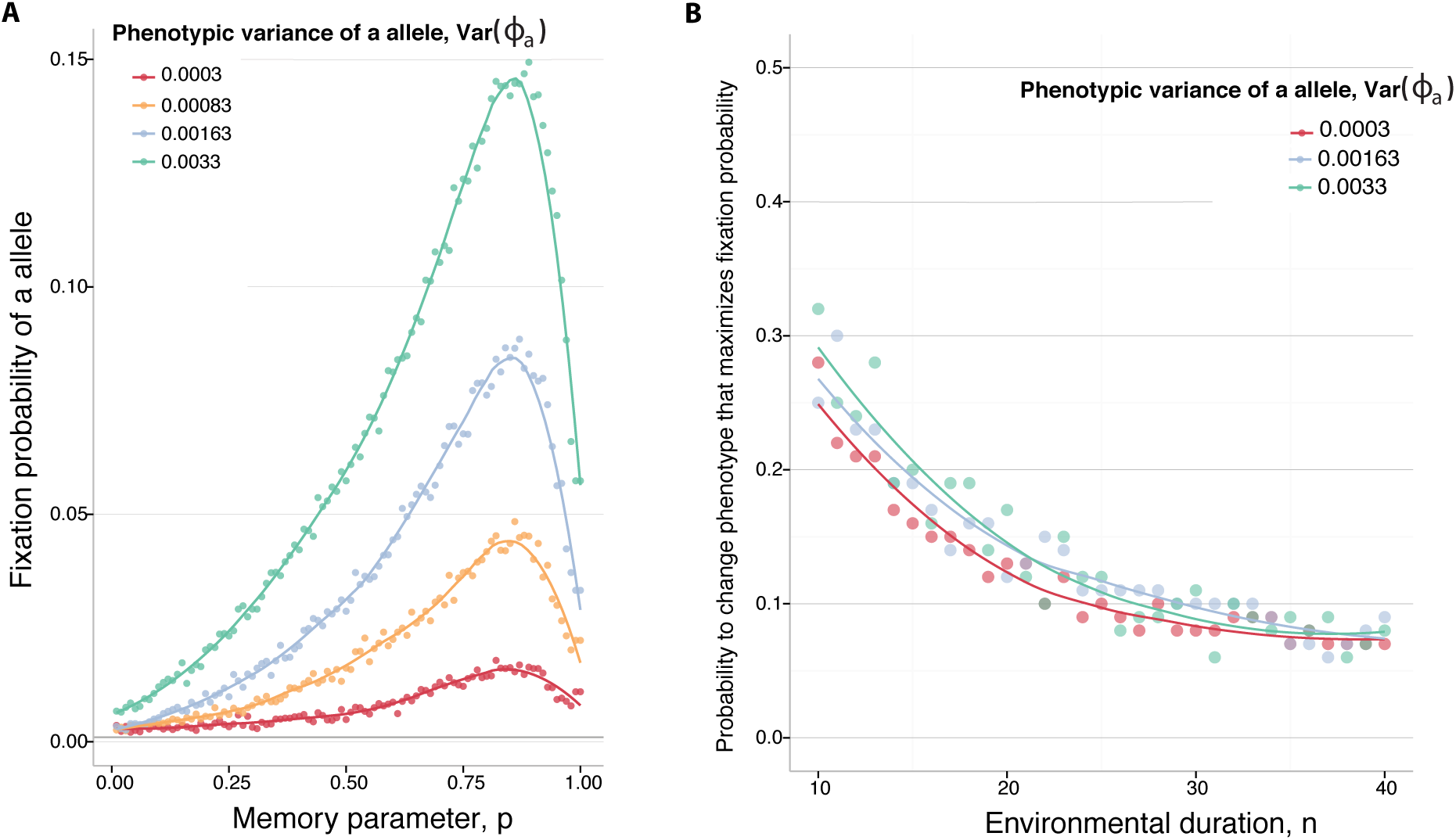
Fixation of a phenotypically variable allele in a periodic environment. **Panel A:** The phenotypic memory *p*, on the *x*-axis, is plotted against the fixation probability of a novel *a* allele in a population of size *N* = 1000. Different colors represent different variances of the *a* phenotype, Var(Φ*_a_*), as indicated. The duration of one environmental period is *n* = 20. **Panel B**: The environmental duration is plotted on the *x*-axis, against the value of phenotypic memory *p\** that maximizes the probability of fixation in a population of size *N* = 1000. Different colors represent different phenotypic variances of the *a* allele, Var(Φ*_a_*), as indicated. The curves show cubic spline fits to the simulated data.

## Discussion

All cell populations harbor phenotypic variation, even cells that are genetically identical and in carefully controlled laboratory environments [7]. This stochastic variation is often regarded as merely experimental noise, and measurements are often averages of this variation [44]. However, transient and variable phenotypes can be mediated by transitions in the epigenome, and they may provide an additional, selectable layer of traits which is increasingly recognized as an evolutionary driving force in many biological systems. From bacterial infections, to the adaptive immune system, or neoplastic development, this form of phenotypic variability can provide a population with increased opportunity to adapt under varying selection pressures [5; 41; 45].

We have studied the fate of an allele that endows the potential to express different, partly heritable phenotypes, in a population that otherwise lacks the capacity for phenotypic heterogeneity. We showed that the fixation probability of such an allele depends critically on the probability of phenotypic inheritance between parent and offspring, in both constant and fluctuating environments. Whereas in constant environments phenotypic memory always increases the fixation probability of a plastic allele, in periodically changing environments, by contrast, there exists an intermediate phenotypic memory that maximizes the invasion probability of the phenotypically plastic allele. In the clinical contexts of infection or tumor spread these results have an immediate qualitative implication: therapeutic interventions that either greatly reduce or greatly increase the heritability of cellular phenotypes/epi-states are expected to reduce the advantage otherwise enjoyed by plastic populations of pathogenic cells (e.g. persister bacterial cells or neoplastic subpopulations that vary in motility).

Previous theoretical studies on the adaptive benefit of phenotypic variability and bet-hedging have mostly been pursued by considering the evolution of (epi)mutation rates in fluctuating environments by modifier loci, in an infinite population. These studies have found that (epi)mutation rates should evolve in tune with the correlation between the environments of parent and offspring [46; 47]. When the environment fluctuates periodically between two states with different optimal phenotypes, the uninvadvable switching rate between phenotypic states will evolve to approximately 1/*n*, where *n* is the number of generations between environmental changes. This un-invadable switching rate is an evolutionary stable strategy (ESS) in an infinite population [48]. Further studies confirmed these results and also generalized them to include both environmental and spatial fluctuations in selection [47; 49; 50]. All these studies, however, consider the dynamics of mutation rates in infinite populations, and they do not explore the evolutionary fate of a new mutation in a finite population subject to demographic stochasticity.

The traditional definition of an ESS in an infinite population is not very useful in finite populations, where even inferior alleles have a positive probability of fixation. As a result, the notion of strict ESS is typically inaccurate for predicting long-term evolutionary dynamics in finite populations [51; 52]. In an infinite population, the long term dynamics of a new mutant do not depend on the initial environment or phenotypes present in the population, but these factors can prove crucial for the early dynamics in a finite population. Fixation probabilities, on the other hand, are much better metrics that can capture the long-term evolutionary dynamics in finite populations [52; 53].

We have focused our analysis on a problem that is similar to the one underlying the ESS – namely, finding the strategies that maximize the fixation probability of a new mutant in a finite population. We have found that these strategies behave similarly to the ESS strategies in infinite populations. This type of concordance between infinite and finite population models is relatively rare in population-genetic settings. Therefore, to further compare our results with those found in infinite populations we have also analyzed the case of a two-phenotype switching model (as illustrated in **Supplementary Figure S5**). Once again, we find an intermediate switching rate that maximizes the fixation probability of a new mutant in a finite population (**Supplementary Figure S6**). We have compared this optimum switching rate with the optimum ESS strategy previously derived in infinite populations [54]. **Supplementary Figure S7** shows that these two optima, found in these two very different frameworks, are remarkably concordant, across a full order of magnitude in population size variation, extensive variation in environmental period, and also variation in phenotypic variance (not shown).

It is also useful to compare our finite-population analysis to results on ESS switching rates in terms of long-term geometric mean fitness. Ishii et al. [46] showed that, in infinite populations, the ESS switching rate maximizes a population’s long-term geometric mean fitness. This result has been generalized in related models of phenotypic switching, also in infinite populations [54]. To compare to these infinite-population results, we analyzed the evolutionary dynamics of a finite population fixed for the *a* allele, and we computed the geometric mean fitness across a full environmental period (that is, across 2*n* generations) of the mean phenotype in the population at each generation, after reaching stationary state. We found that this stationary geometric mean fitness is, again, a non-monotonic function of the phenotypic memory parameter *p* (**Figure 5, Panel A**). Moreover, the memory *p\** that maximizes the stationary geometric mean fitness also maximizes the fixation probability of a novel *a* mutant in the population (**Figure 5, Panel B**). The coincidence of the phenotypic memory that maximizes fixation probability and the memory that maximizes stationary geometric mean fitness helps to explain the strong concordance between our finite-population results and the prior literature in infinite populations, where the relationship between ESS and geometric mean fitness was already established [46; 54].

**Figure 5:**
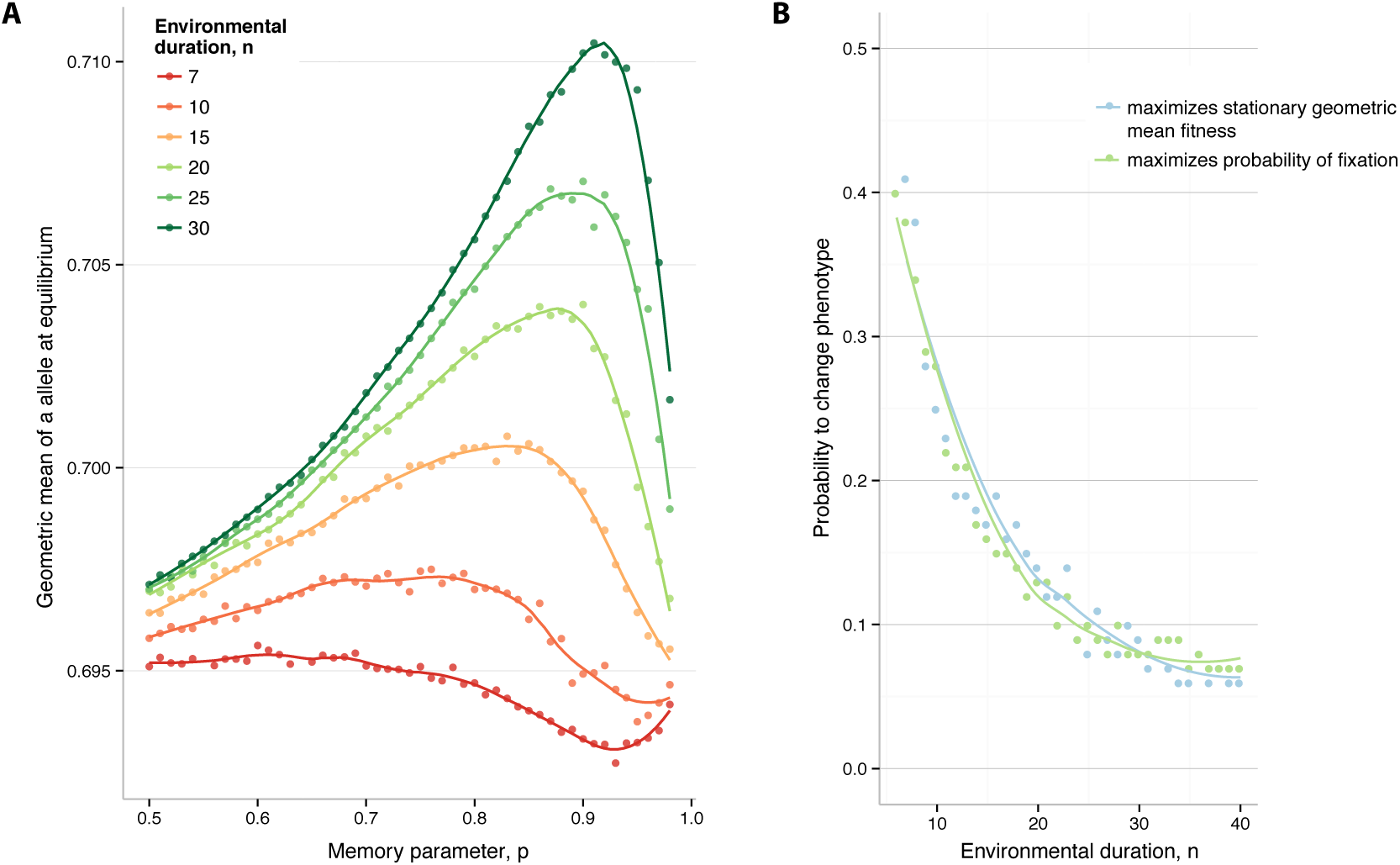
Fixation probabilty and geometric mean fitness. **Panel A:** Geometric mean fitness in stationary state, as a function of phenotypic memory *p*. The colors represent different environmental durations *n*, as indicated. Population size *N* = 1000. Variance of the *a* phenotype, Var(Φ*_a_*) = 0.0133. **Panel B:** Comparison of the phenotypic memory that maximizes the fixation probability and the memory that maximizes stationary geometric mean fitness. Population size *N* = 1000. Variance of the *a* phenotype, Var(Φ*_a_*) = 0.0133. The curves show cubic spline fits to the simulated data.

The goal of this study has been to develop a population-genetic model of the effects of bet-hedging in populations that harbor (partly) heritable variation in expressed phenotypes. Although this study has been largely theoretical, and it neglects the myriad mechanisms of phenotypic plasticity that arise in, and differ among, biological systems, the models we have developed nonetheless provides a logical basis for qualitative and, eventually, even quantitative predictions in biological systems. Such predictions are increasingly understood as required for rational design of therapies in clinical contexts, where phenotypic heterogeneity and epi-states inducing quiescent cellular states may be the primary driver of resistance and therapy failure across a diverse array of diseases [11; 55]. There is a need to consider the selective pressures on alleles that endow cellular populations phenotypic plasticity. Our results provide insight into how, and under what conditions, such phenotypic heterogeneity can provide an initial selective advantage in changing environmental regimes.

Our results suggest that there exist two very different, almost opposing, types of intervention in diseases characterized by cellular populations with plastic phenotypes. Both intervention options are based on the fact that, unlike genetic changes, such as mutations and chromosomal rearrangements, expression or phenotypic changes are rapidly reversible. The existence of an intermediate phenotypic heritability that maximizes the fixation probability of the plastic allele suggests an effective intervention by disrupting the molecular memory to either extreme (memory *p* = 0 or *p* = 1), e.g. by disrupting key signaling pathways that regulate epi-states. In the context of several cancer types, for example, tumor imaging has shown that a small proportion of cancer cells are motile (around 15%, [17; 56]) and able to overcome inhibitors that target a particular mode of movement by switching their motility phenotype. To curtail the invasion and spread of these cells, one strategy could be to manipulate the phenotypic memory of such variants, e.g. by blockage of RAC signaling pathways, to prevent phenotypic conversion between rounded amoeboid cells and elongated cells (i.e. increasing phenotypic memory). Conversely, inhibition of ROCK-family kinases leads to RAC reactivation, thereby reducing phenotypic memory [17; 57; 58]. Therapies that target either RAC or ROCK function may therefore reduce the resistance advantage that cancer cell populations otherwise enjoy in the face of selective therapy. Of course, further predictive models based on specific pathways, and unique to particular cellular populations, are needed to formulate optimal therapies that minimize the spread of phenotypically-heterogeneous genotypes and resistance. And the most efficacious ways to implement these strategies remain to be defined.

---

**Figure S1.**
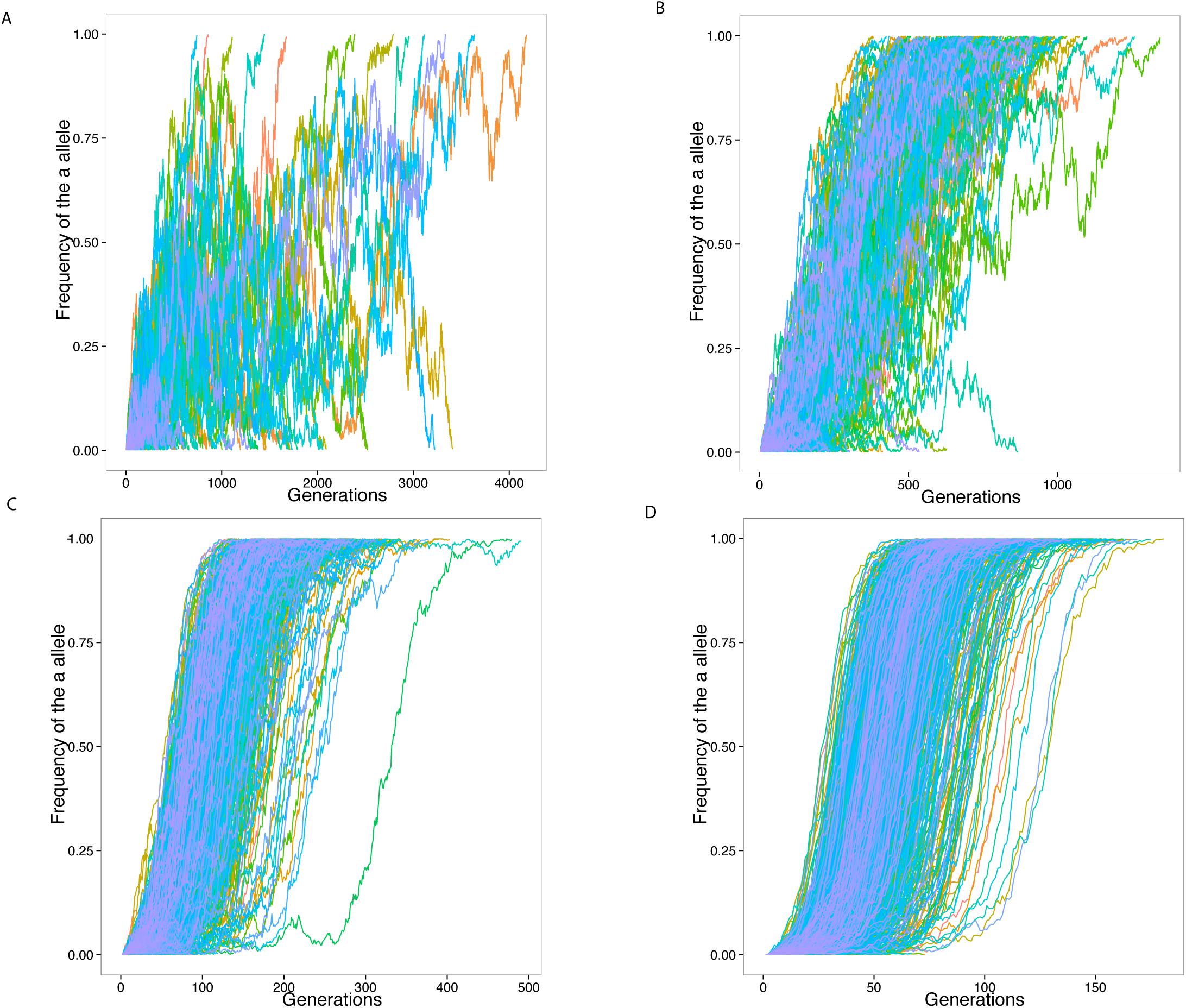
Example sample paths for 10,000 runs of the simulation in a constant environment. The phenotype of *A* allele fixed at Φ*_A_* = *δ*_0_._8_. Mean phenotype of *a* allele is also 0.8, 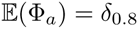, with variance Var(Φ*_a_*) = 0.0133. The y-axis shows the frequency of the *a* allele in the population. Population size *N* = 1000. **Panel A:** Memory parameter *p* = 0. **Panel B:** Memory parameter *p* = 0.3. **Panel C:** Memory parameter *p* = 0.7. **Panel D:** Memory parameter *p* = 0.9.

**Figure S2.**
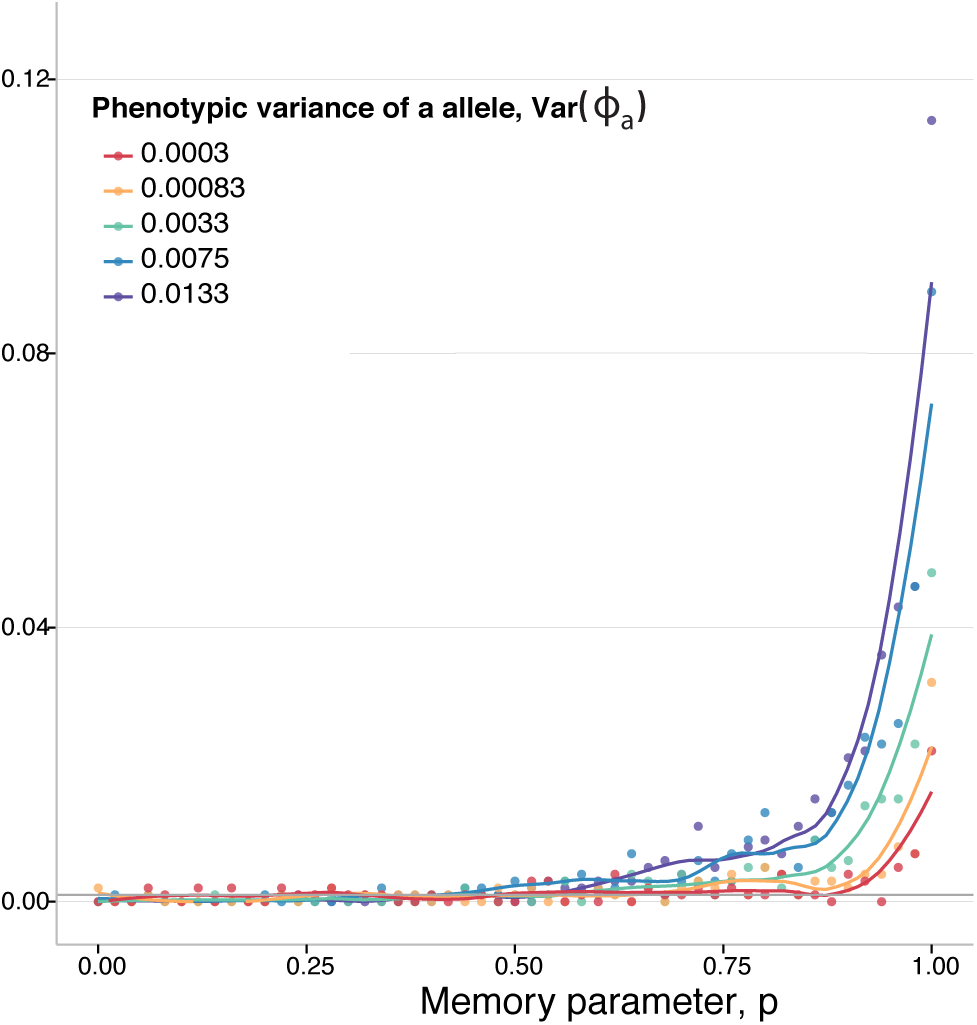
The impact of global versus individual-specific memory in a constant environment. A model where, every generation, all the individuals of *a* genotype draw the same phenotype. Moreover, at each generation, the probability of phenotypic memory *p* is globally determined for all individuals of *a* genotype. Other parameters are the same as **Figure 2**. The curves represent a fit to the data using a generalized additive model with penalized cubic regression splines.

**Figure S3.**
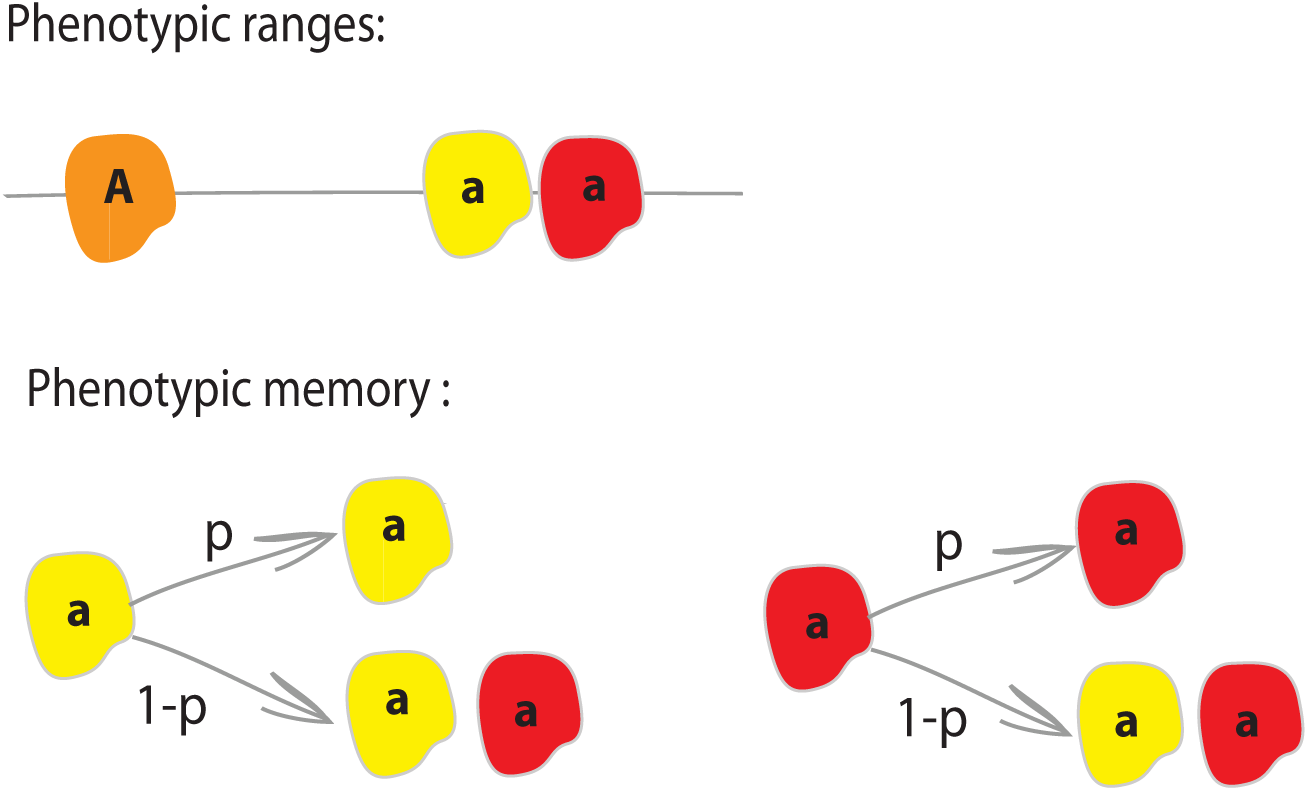
Illustration of the constant environment model with discrete distribution. The major differentiator of this simplified model is the fact that the *a* allele only has access to two discrete phenotypes, one with higher fitness than the *A* allele, and one with lower fitness.

**Figure S4.**
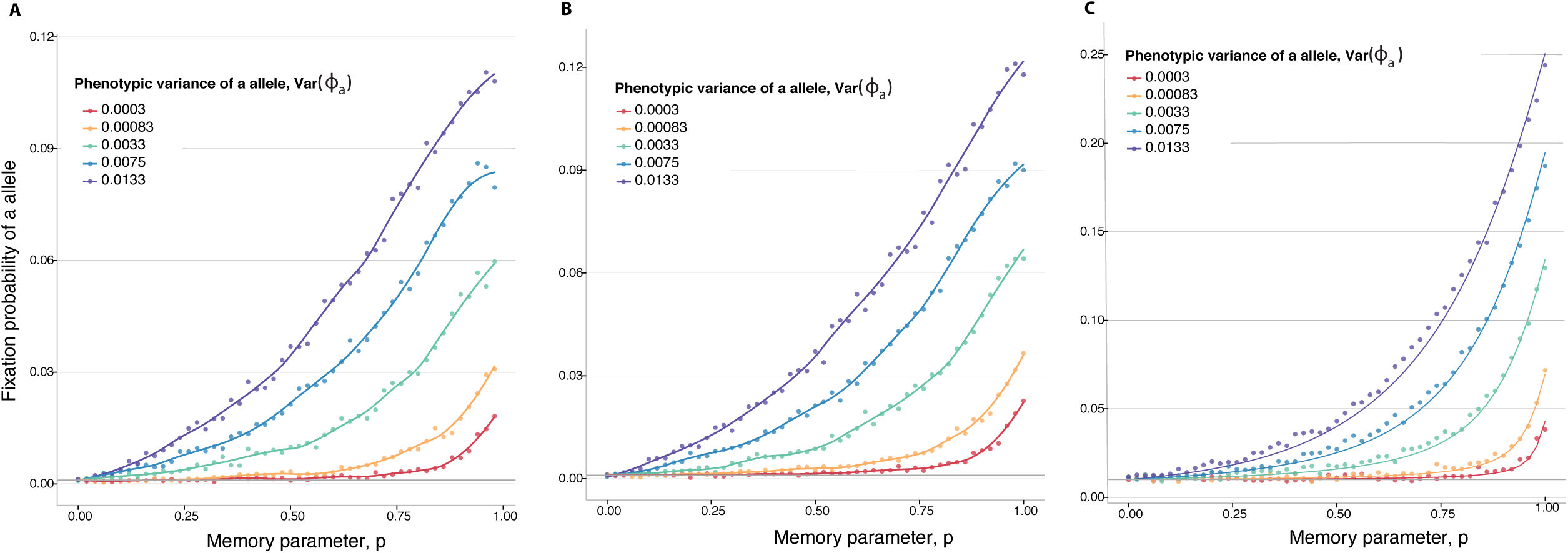
Invasion probabilities in a constant environment. **Panel A:** Continuous phenotypic distribution of the *a* allele and is the same as **Figure 2**. The curves represent a fit to the data using a generalized additive model with penalized cubic regression splines. **Panel B**: Equivalent discrete phenotypic distribution for *a,* as illustrated in **Figure S3**. All parameters are the same as **Panel A**. The curves represent a fit to the data using a generalized additive model with penalized cubic regression splines. **Panel C**: Comparison of simulations and the analytical approximation using the discrete model. We assume the first copy of the *a* allele is always the one with higher fitness, hence there is an increase in the probabilities of fixation. The dots in this panel represent the result of the simulation, while the curves represent the analytical approximations.

**Figure S5.**
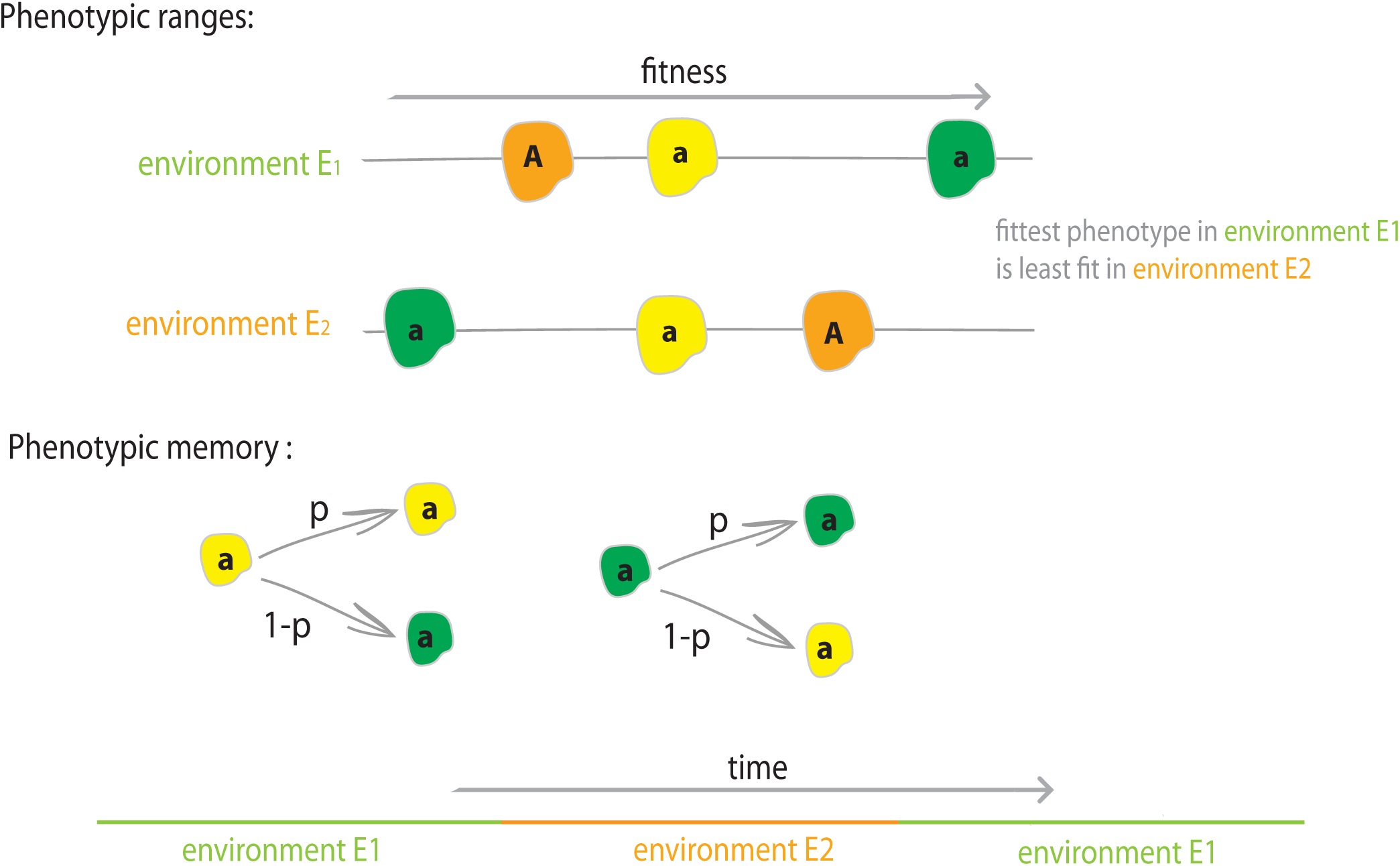
Illustration of the periodic environment model with discrete distribution and phenotypic switching. The major differentiator of this simplified model is the fact that the *a* allele only has access to two discrete phenotypes. Moreover, *p* in this model represents the probability of phenotypic switching, whereby the new *a* phenotype is no longer resampled from the entire distribution, but is instead switched to the alternative phenotypic state.

**Figure S6.**
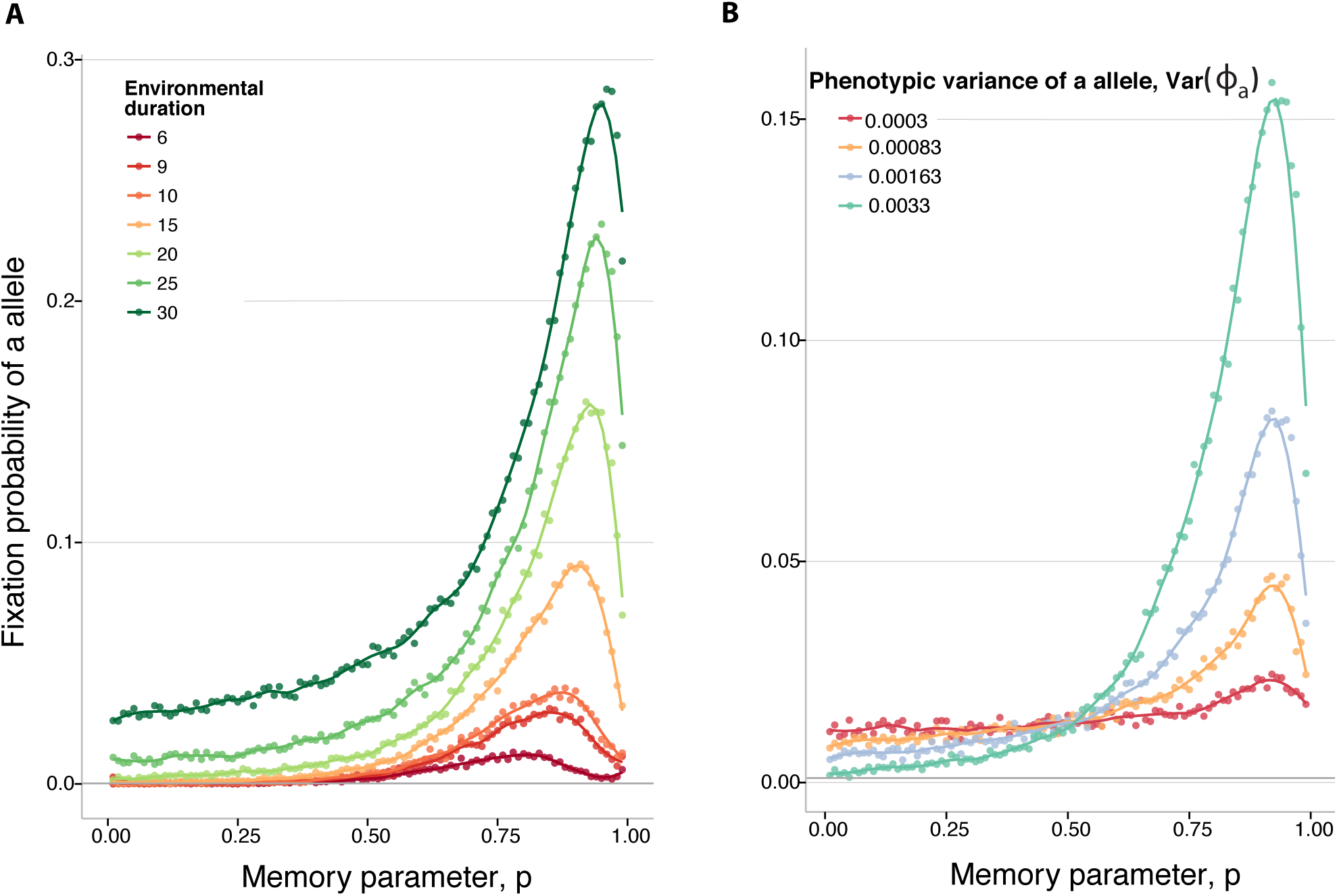
The phenotypic switching model. 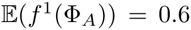 and 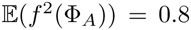 The initial environment is *E*_1_. Population size *N* = 5000. **Panel A**: The colors represent different rates of environmental change, *n*, as presented in the legend. Fitness of a allele sampled between two different phenotypes such that the binomial variance is adjusted to be equal to the equivalent uniform continuous distribution presented in **Figure 2**. **Panel B**: The duration of one environmental stretch is equal to 20 generations. The colors represent different binomial variances of the *a* allele, Var(Φ*_a_*), as presented in the legend. The curves represent a fit to the data using a generalized additive model with penalized cubic regression splines.

**Figure S7.**
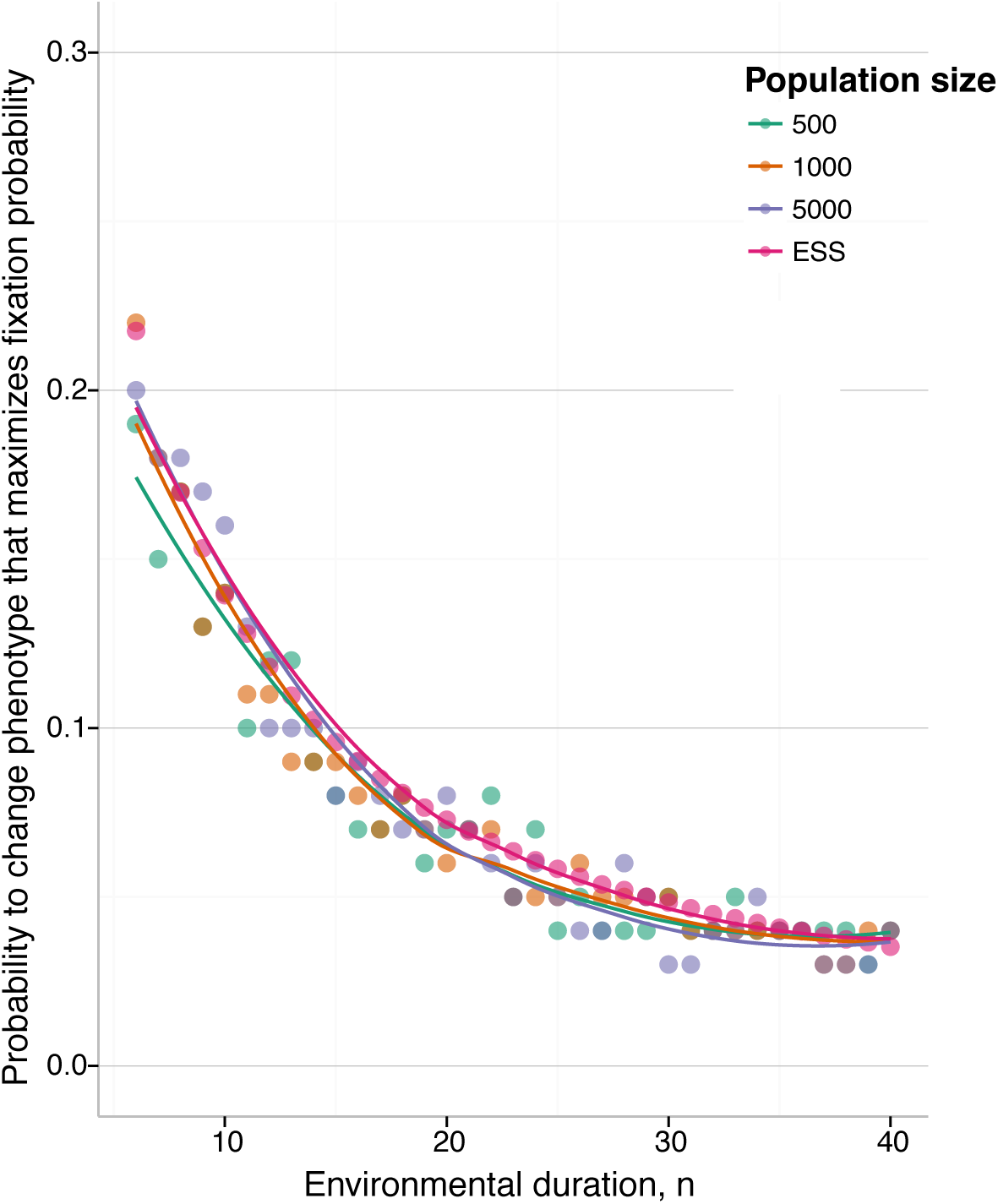
Phenotypic memory that maximizes the fixation probability with phenotypic switching. The y-axis shows the phenotypic memory that maximizes the probability of fixation, for different population sizes *N*. Variance of the *a* phenotype is Var(Φ*_a_*) = 0.0133. The ESS switching rates found in the infinite population case are also plotted. The curves represent a fit to the data using a generalized additive model with penalized cubic regression splines.

